# Design an efficient multi-epitope peptide vaccine candidate against SARS-CoV-2: An in silico analysis

**DOI:** 10.1101/2020.04.20.051557

**Authors:** Zahra Yazdani, Alireza Rafiei, Mohammadreza Yazdani, Reza Valadan

**Affiliations:** Department of Immunology, Molecular and Cell Biology Research Center, School of Medicine, Mazandaran University of Medical Sciences, Sari, Iran; Department of Chemistry, Isfahan University of Technology, Isfahan, 84156-83111, Iran

**Keywords:** SARS-CoV-2, multi-epitope vaccine, structural proteins, humoral immunity, cellular immunity

## Abstract

**Background:** To date, no specific vaccine or drug has been proven to be effective for SARS-CoV-2 infection. Therefore, we implemented immunoinformatics approach to design an efficient multi-epitopes vaccine against SARS-CoV-2.

**Results:** The designed vaccine construct has several immunodominant epitopes from structural proteins of Spike, Nucleocapsid, Membrane and Envelope. These peptides promote cellular and humoral immunity and Interferon gamma responses. In addition, these epitopes have antigenicity ability and no allergenicity probability. To enhance the vaccine immunogenicity, we used three potent adjuvants; Flagellin, a driven peptide from high mobility group box 1 as HP-91 and human beta defensin 3 protein. The physicochemical and immunological properties of the vaccine structure were evaluated. Tertiary structure of the vaccine protein was predicted and refined by I-Tasser and galaxi refine and validated using Rampage and ERRAT. Results of Ellipro showed 242 residues from vaccine might be conformational B cell epitopes. Docking of vaccine with Toll-Like Receptors 3, 5 and 8 proved an appropriate interaction between the vaccine and receptor proteins. In silico cloning demonstrated that the vaccine can be efficiently expressed in *Escherichia coli.*

**Conclusions:** The designed multi epitope vaccine is potentially antigenic in nature and has the ability to induce humoral and cellular immune responses against SARS-CoV-2. This vaccine can interact appropriately with the TLR3, 5, and 8. Also, this vaccine has high quality structure and suitable characteristics such as high stability and potential for expression in *Escherichia coli*.

## 1. Introduction

Severe acute respiratory syndrome coronavirus 2 (SARS-CoV-2) which the cause of COVID-19 disease, was first reported as a pneumonia epidemic in the Chinese city of Wuhan (Hubei province) in December 31, 2019 belongs to the Beta coronavirus genus[1-3]. Coronaviruses are positive-sense single-stranded RNA viruses. They belong to the order of Nidovirales and superfamily of orthocoronaviridae. This superfamily has four genus including: alpha, beta, gamma and delta coronaviruses. Gamma and delta coronaviruses infect birds, while alpha and beta coronaviruses generally infect mammals such as human. In immunocompetent individuals, they generally cause mild respiratory infections, such as common cold, while in some humans coronavirus infections cause serious disease, such as SARS (Severe Acute Respiratory Syndrome) and MERS (Middle East Respiratory Syndrome) epidemics.

The SARS-CoV-2, the viral cause of COVID-19, has a large mRNA genome, 26 to 32 kb in length, meaning a 5’ cap cover structure and a 3’ polyadenylated. This encodes several structural and nonstructural proteins. Among structural proteins, spike (S), envelope (E), membrane (M), and nucleocapsid (N) proteins are important in inducing immune responses [3-5].Protein S is the main tool for virus entry into the cell, which interacts with host cell receptor, angiotensin-converting enzyme 2 (ACE2). ACE2 is a metallopeptidase expressed in variant tissues, including alveolar epithelial cells, fibroblasts, endothelial cells, and enterocytes. [6-10].The N protein is an essential RNA-binding protein for transcription and replication of the viral RNA. It has the significant roles in forming of the helical ribonucleoproteins during packaging the RNA genome, regulating of viral RNA synthesis during replication, transcription and modulating of metabolism in the infected cells [11-14]. T cell responses against the S and N proteins of SARS-CoV virus are most dominant and long-lived than other structural proteins[15]. protein E has an important role in the assembly of the viral genome[16, 17].The M protein plays a pivotal role in virus assembly,budding and replication of virus particles in the host cells[18].

SARS-CoV-2 is transmitted quickly and causes considerable fatality rate, so that World Health Organization (WHO) reported over 2 114 269 confirmed cases globally and at least 145 144 deaths because of this disease until April 17, 2020[19]. There is currently no vaccine or approved treatment for this disease. One of the major challenges for scientists in dealing with the new coronavirus pandemic will be to get a useful and effective vaccine. For this reason, trying to develop an effective vaccine to control the virus has been the subject of research by many scientists around the world [4, 20-22]. However, achieving such a vaccine with conventional methods is very time consuming and expensive. As such, *in silico* prediction of the vaccine targets is very important because the targets can be selected with a more open eye in a limited time and resources. Bioinformatic approaches are very helpful to identify the effective epitopes and developing vaccine. Several vaccines for virus diseases were designed based on these approaches that include effective vaccines on Human papilloma viridea (HPV) [23],Ebola [24], Zika[25]and MERS[26, 27]viruses. There are also few reports of a vaccine candidate for COVID19[28, 29]. These vaccines contained multiple cytotoxic T lymphocytes (CTL) and B cell epitopes against several proteins of SARS-CoV-2 virus. However, those vaccines didn’t protect against all structural immunodominant proteins of virus. Therefore, to overwhelm these limitations, we designed a multi-epitope peptide vaccine consisted of S, M, N, and E viral proteins. The vaccine has appropriate physicochemical properties such as stability at room temperature, antigenic capability, and no possibility of allergy. This multi-epitope vaccine consists of three epitopes from S protein and one epitope from each of M, N, and E proteins, respectively. Each of the epitopes has a high ability to stimulate humoral and cellular immune responses, especially the proper production of Interferon gamma (IFN-γ). In addition, we used appropriate adjuvants in the vaccine structure to potentiate the immunogenicity of the antigens.

Therefore, we incorporated the potent adjuvants of N and C terminus of Flagellin as a toll like receptor (TLR) 5, a driven peptide from high mobility group box1 (HMGB1) as HP-91 and human beta defensin 3 (HBD-3) in the construct of the vaccine. The vaccine segments were connected to each other by appropriate linkers. Physicochemical properties, structural stability, and immunological characterizations of the vaccine were evaluated. Modeling, refinement, and validation were performed to access a high quality three dimensional (3D) structure of the vaccine protein. Docking evaluation showed an appropriate interaction between the vaccine and TLRs 3, 5 and 8. In silico cloning showed that the vaccine could be effectively expressed in *E.coli*. Totally, a potential vaccine candidate with proper immunological and stable physicochemical properties against SARS-CoV-2 was designed. It is expected the vaccine could be capable to protect humans from COVID19 disease.

## 2. Methodology

### 2.1 Protein sequence retrieval

The protein sequences of spike glycoprotein (accession number of QIC53213.1), nucleocapsid phosphoprotein (QHU79211.1), membrane glycoprotein (QIC53207.1) and envelope protein (QIC53215.1) were driven using NCBI (https://www.ncbi.nlm.nih.gov/) databases and saved in FASTA format for subsequent analysis.

### 2.2 Screening of potential epitopes

#### 2.2.1 Screening of Major histocompatibility complex class I (MHC-I) epitopes

MHC-I humans alleles with 9 mer length were selected using IEDB (www.iedb.org) database through IEDB recommended 2.22 method and Net-MHC 4.0 online server (http://www.cbs.dtu.dk/services/NetMHC/).IEDB (Instructor/Evaluator Database) is a repository of web-based tools which predicts and analysis of immune epitopes. It uses a consensus method based on artificial neural network (ANN), stabilized matrix method (SMM), and Combinatorial Peptide Libraries (CombLib), if any corresponding predictor is available for the peptide sequence [30-48]. NetMHC 4.0 software is a free server for the prediction of peptide-MHCI binding affinity by gapped sequence alignment. Prediction methods of this server are based on alignments that include insertions and deletions and have significantly higher performance than methods based on peptides of single lengths methods[32].

#### 2.2.2 Screening of Cytotoxic T lymphocytes (CTL) epitopes

Prediction of CTL epitopes was done by an online server CTL Pred. This method is based on quantitative matrix (QM) and machine learning techniques for example, artificial neural network (ANN) and support vector machine (SVM).The server also uses the consensus and combined prediction approaches. The consensus and combined prediction approaches are more specific and sensitive, respectively, in comparison with individual approaches such as ANN and SVM[49].

#### 2.2.3 Screening of Human leukocyte antigen (HLA)-II epitopes

IEDB database (IEDB recommended 2.22 method) [40, 50-53] (www.iedb.org) and RANKPEP online server (http://imed.med.ucm.es/Tools/rankpep.html) were employed for screening of HLA-II epitope. The database of IEDB was described previously. The RANKPEP server predicts MHC-II binding epitope by position specific scoring matrices (PSSMs) that are structurally consistent with the binding mode of MHC-II ligands [54].

#### 2.2.4 Screening of Linear B cell epitopes

Prediction of linear B cell epitope was done by IEDB[55] (Emini surface accessibility method) and BepiPred software. BepiPred 2.0 predicts the location of linear B-cell epitopes by a combination of a hidden Markov model and a propensity scale method. The epitope Threshold of this server was selected 0.5[56].

### 2.3 Selection of the epitope segments

The results of all the above predictions were pooled and compared together, and the regions with the highest overlaps were determined. These immunodominant regions were employed for future analyses to finally select the most appropriate epitope domains for vaccine construct.

### 2.4 In silico analyzing of IFN-γ inducing epitopes

IFN epitope server (http://crdd.osdd.net/raghava/ifnepitope) was employed for determining the induce IFN-γ production ability in the selected epitopes. This web server classifies MHC binder epitopes into IFN-γ inducing (positive numbers) and non-inducing IFN-γ (negative numbers) using several methods such as; motifs-based search, machine learning technique and a hybrid approach. Accuracy of Best prediction based on hybrid approach in this software is 82.10%[57].

### 2.5 Evaluation of antigenicity

Antigencity of epitopes was assessed by ANTIGENpro (http://scratch.proteomics.ics.uci.edu/) and VaxiJen v2.0 (http://www.ddg-pharmfac.net/vaxijen/VaxiJen/VaxiJen.html) servers. Web server of ANTIGENpro is the first sequence-based, alignment-free, and pathogen independent predictor, using protein antigenicity microarray data for predicting the protein antigenicity[58]. VaxiJen is also the first server for prediction of antigens. This server applies a new alignment-independent approach that is based on auto-cross covariance (ACC) transformation of protein sequences into uniform vectors of principal amino acid properties. Depending on the target organism (bacteria, virus or tumor) accuracy of this server varies from 70% to 89%. The Threshold of this server was selected 0.5[59, 60].

### 2.6 Allergenicity assessment of predicted epitopes

For predicting protein allergenicity with a high accuracy, the software of AllergenFP v.1.0 (http://ddg-pharmfac.net/AllergenFP/) was used. This is an online server that identifies allergens based on amino acid principle properties such as hydrophobicity, helix and β-strand forming with accuracy about 88% [40].

### 2.7 Vaccine engineering, evaluation of physicochemical properties, antigenicity and allergenicity

According to the results of the previous steps, four epitopes from S protein, two epitope from N protein and one epitope from E protein were selected to be incorporated in the vaccine construction. The whole construction was designed using joining these epitopes to two adjuvant sequences of N and C terminals of Flagellin and the synthesis peptide of HP-91.Various segments of the designed-vaccine connected to each other by suitable linkers.

After designing the vaccine, Several physicochemical parameters including molecular weight, theoretical isoelectric point (pI), total number of positive and negative residues, extinction coefficient,

Instability index, half-life, aliphatic index, and grand average hydropathy (GRAVY) were computed using Expasy’s ProtParam at http://us.expasy.org/tools/protparam.html [61]. Antigenicity of the final vaccine construct was evaluated using vaxijen v2.0 and ANTIGENpro and allergenicity was assayed by AllergenFP v.1.0 server.

### 2.8 Tertiary structure development

The 3-D modeling of designed construct was predicted using the I-TASSER software (https://zhanglab.ccmb.med.umich.edu/I-TASSER/) [61].This server showed five models as a result of the prediction. This model was accepted as a vaccine structure with the highest reliability score (c-sore) and was refined using Galaxy Refine web services. It successfully tested in CASP10 (Critical assessment of techniques for protein structure prediction) experiments. This server refines the whole protein with gentle and aggressive relaxation methods. This first reconstructs all side-chain conformations and repeatedly relaxes the structure with short molecular dynamics simulations after side-chain repacking perturbations [62].Finally, Galaxy Refine showed five structures as refined models of vaccine structure. The RAMPAGE (http://mordred.bioc.cam.ac.uk/~rapper/rampage.php), and ERRAT (http://services.mbi.ucla.edu/ERRAT/) servers were used to validate the refined 3D structures obtained from Galaxy Refine web service. [63, 64]. Finally, the refined and high-quality 3D structure of the vaccine was observed by PyMOL software v2.1.1.PyMOL is an open-source that widely used for bimolecular function[65].

### 2.9 Prediction of discontinuous B cell epitope

Discontinuous B-cell epitopes were predicted from the 3D vaccine structure using Ellipro in IEDB database (http://tools.immuneepitope.org/tools/ElliPro).ElliPro is a useful research tool for identifying immunoepitopes in protein antigens which implements Thornton’s method and with a residue clustering algorithm.The MODELLER program and the Jmol viewer allow the prediction and visualization of immunoepitopes in a given protein sequence or structure. In comparison with six other structure-based methods that are using for epitope prediction, ElliPro performs the best prediction and gave an AUC value of 0.732, when the most significant prediction is considered for each protein. [66].

### 2.10 Molecular docking with TLR-3, 5 and 8

The first, tertiary structure of the human TLR-3, 5, and 8 was obtained from PDB database (www.rcsb.org) with codes of 3J0A, 5GS0 and 3w3g, respectively. Next, protein-protein docking of the vaccine structure (as ligand) and each TLR (as receptor) was performed by CLUSPRO 2.0 online server (cluspro.bu.edu/login.php). CLUSPRO 2.0 is a fully automated and fast rigid-body protein–protein docking server. This server evaluates docking of two interacting proteins based on three techniques: first, the Fast Fourier Transform (FFT) correlation approach, second, clustering of the best energy conformations, third, refining the obtained model using short Monte Carlo simulations and the medium-range optimization method SDU [67-69].

### 2.11 Codon optimization and *in silico* cloning

The reverse translation of the designed-vaccine gene sequence was performed by Sequence Manipulation Suite (https://www.bioinformatics.org/sms2/rev_trans.html) to prepare a suitable vaccine sequence for cloning and expression in an appropriate expression system. The properties of sequence gene such as Codon Adaptation Index (CAI), GC content, and Codon Frequency and Distribution (CFD) have the key roles in attaining a high-level of protein expression in the host were evaluated using GenScript online server (https://www.genscript.com/tools/rare-codon-analysis)[24]. Finally, the restriction sites of XhoI and EcoRV were respectively added to the N- and C-terminus of the vaccine DNA sequence using CLC Sequence viewer v8.0 (http://www.cacbio.com) to facilitate the cloning in *E. coli* expression system.

## 3. Results

### 3.1. Immunoinformatics analysis

Protein S, M, N, and E were used to predict HLA-I binding epitopes (HLA-A, B, and C) using IEDB and NetMHC 4. CTL Pred server was used to predict CTL epitopes of SARS-CoV-2 structural proteins. HLA-II binding epitope (DP, DQ, and DR) from these proteins were predicted using RANKEP and IEDB servers. Prediction of linear B cell epitopes was performed by BepiPred and IEDB. Finally, the obtained epitopes from the comparison of above analyses were applied to future selection (Table.1).

**Table.1.**
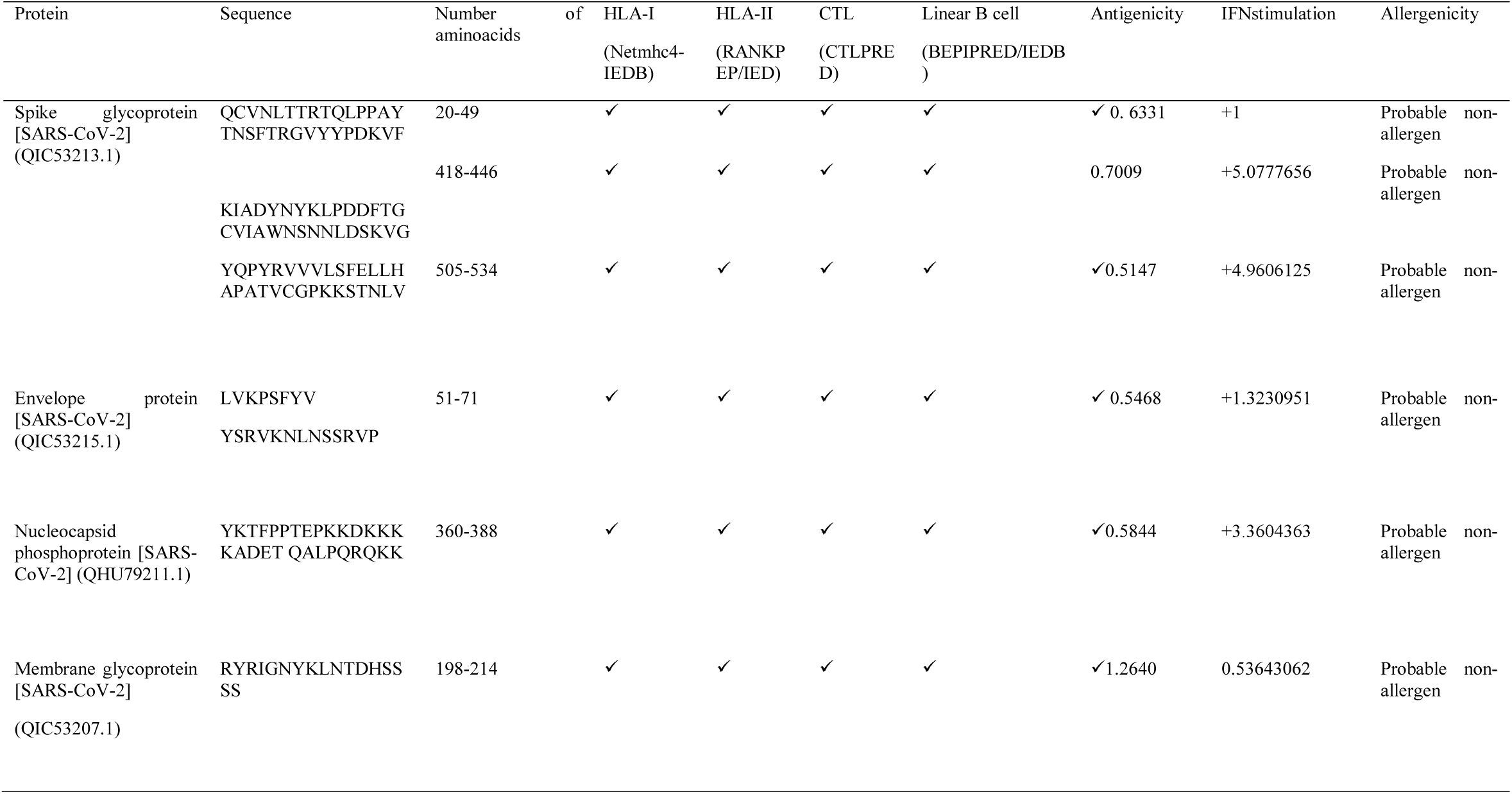
The final immunodominant epitopes were selected from S, M, N, and E proteins of SARS-CoV-2

### 3.2. Antigenic protein screening and avoid allerginicity

All selected peptides were submitted to the IFNepitope server for evaluation their ability to induce IFN-γ. Allergenicity probability of epitopes evaluated by AllergenFPE valuation of epitope antigenicity was performed by ANTIGEN pro and VaxiJen v2.0.The six epitopes (three from S and one from each of E, M, N proteins) that had the highest ability in IFN-γ induction, low allergenicity and potent antigenic ability were chosen to be used in the vaccine structure (Table 1).

### 3.3 Designing a multi-epitopes vaccine construct

Six epitopes with high score which selected to be incorporated in the final vaccine construct were the sequences of 20-49, 418-446, and 505-534 from S, 51-71 from E, 360-388 from N and 198-214 from M protein. In addition, three immunopotent adjuvants Flagellin, HP-91, and HBD-3 were also added to the vaccine structure. The vaccine pieces were linked to each other by using a short linker sequence of LE dipeptide repeats. The final multi epitope peptide vaccine was 473 amino acid residues (Fig.1).

**Figure 1.**
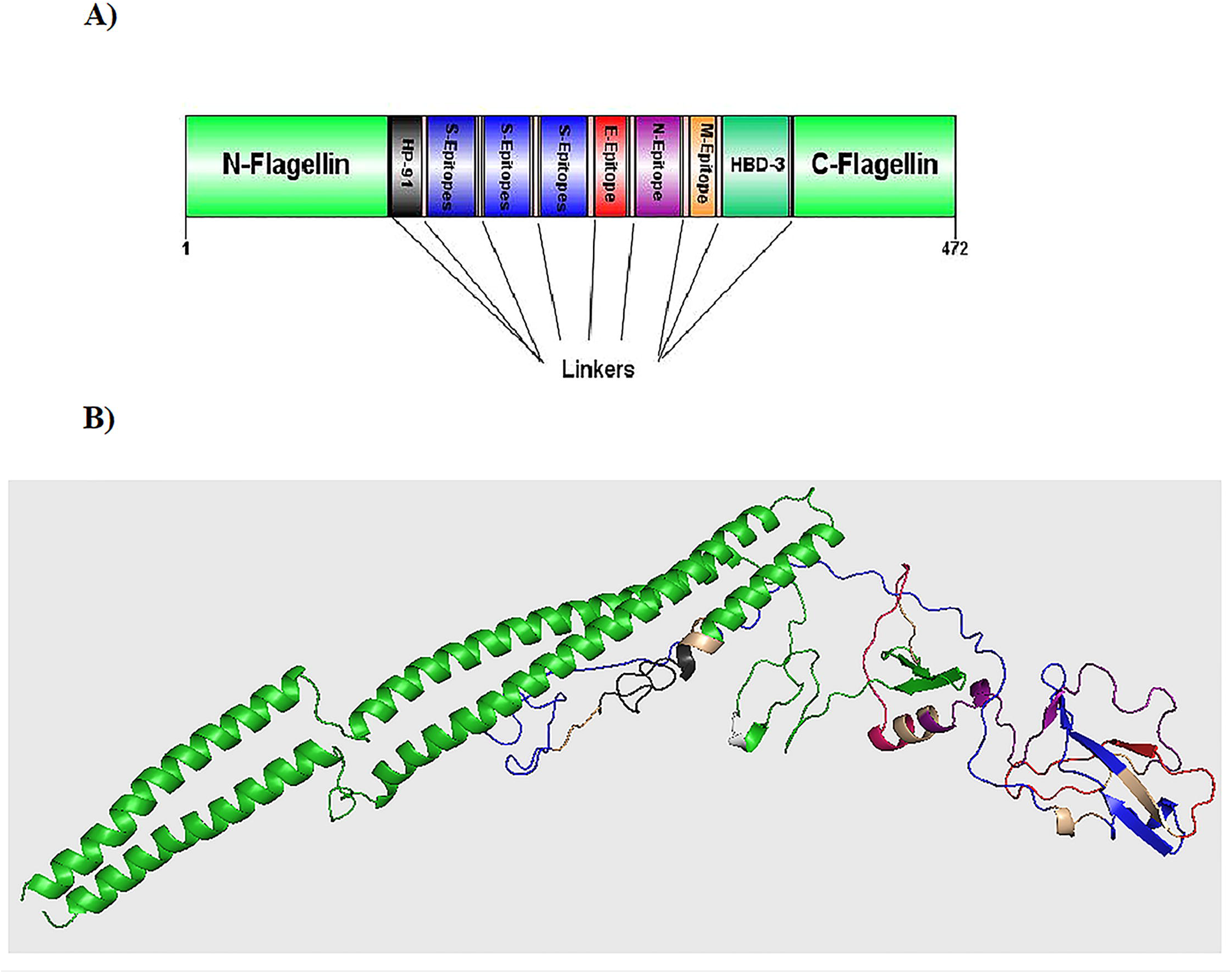
Schematic representation of the designed multi-epitope peptide based-vaccine. The vaccine consist of ten parts: Epitopes from structural proteins S, E, N, and M, adjuvants: Flagellin (in N-and C-terminus), HP-91 and HBD-3 that join to each other by linkers of repeat sequence of LE (A figure). Tertiary structure of the modeled multi-epitope vaccine construct (figure B). The 3D structure of the designed vaccine was predicted via homology modeling by I-Tasser, then the best predicted model was refined by Galaxy Refine and visualized using Pymol software. N-and C-terminus of Flagellinis shown in green, HP-91 in black, S epitopes in blue, E epitopes in purple, N epitope in orange, M epitope in yellow, HBD-3 in green, and linkers are shown in light pink.

### 4.3 Immunological and physicochemical properties of the vaccine structure

To evaluate antigenicity of the whole protein vaccine, the ‘‘virus’’ option was chosen as a target organism. The antigenicity of 0.5319 (probable antigen) was estimated at 0.5% threshold for the virus model. Assessment of antigenicity by Antigen pro showed this vaccine is high antigenic with antigenicity of 0.941411. AllergenFP showed that the T cell epitomes in the vaccine protein are non-allergen. The molecular weight (MW) and theoretical isoelectric point (pI) of the vaccine protein were computed as 51.916kDa and 7.45, respectively. The predicted half-lives were calculated as 30 hour (h) (mammalian reticulocytes, in vitro), 20 h (*yeast*, in vivo), and 10 h (*E. coli*, in vivo). The instability index was 35.77, indicating that the protein vaccine was stable enough (Table.2)

**Table.2.** Analysis of the physicochemical and immunological properties of the SARS-CoV-2 vaccine

| Physicochemical properties | Value |
| --- | --- |
| Molecular Weight (Da) | 51916.54 |
| Instability index | 35.77 |
| Gravy | -0.458 |
| Aliphatic index | 87.03 |
| Theoretical pI | 7.45 |
| No. amino acids | 472 |
| Total no. of negatively charged residues<br>(Asp+Glu) | 53 |
| Total no. of positively charged residues<br>(Arg+Lys) | 54 |
| No. of atoms | 7308 |
| Antigenicity |  |
| Antigenicity / ANTIGENpro | 0.941411 |
| Antigenicity /vaxijen | 0.5319 ( Probable antigen ) |
| Allergenicity / AllergenFP v.1.0 | Probable non-allergen |

**Table.3:**
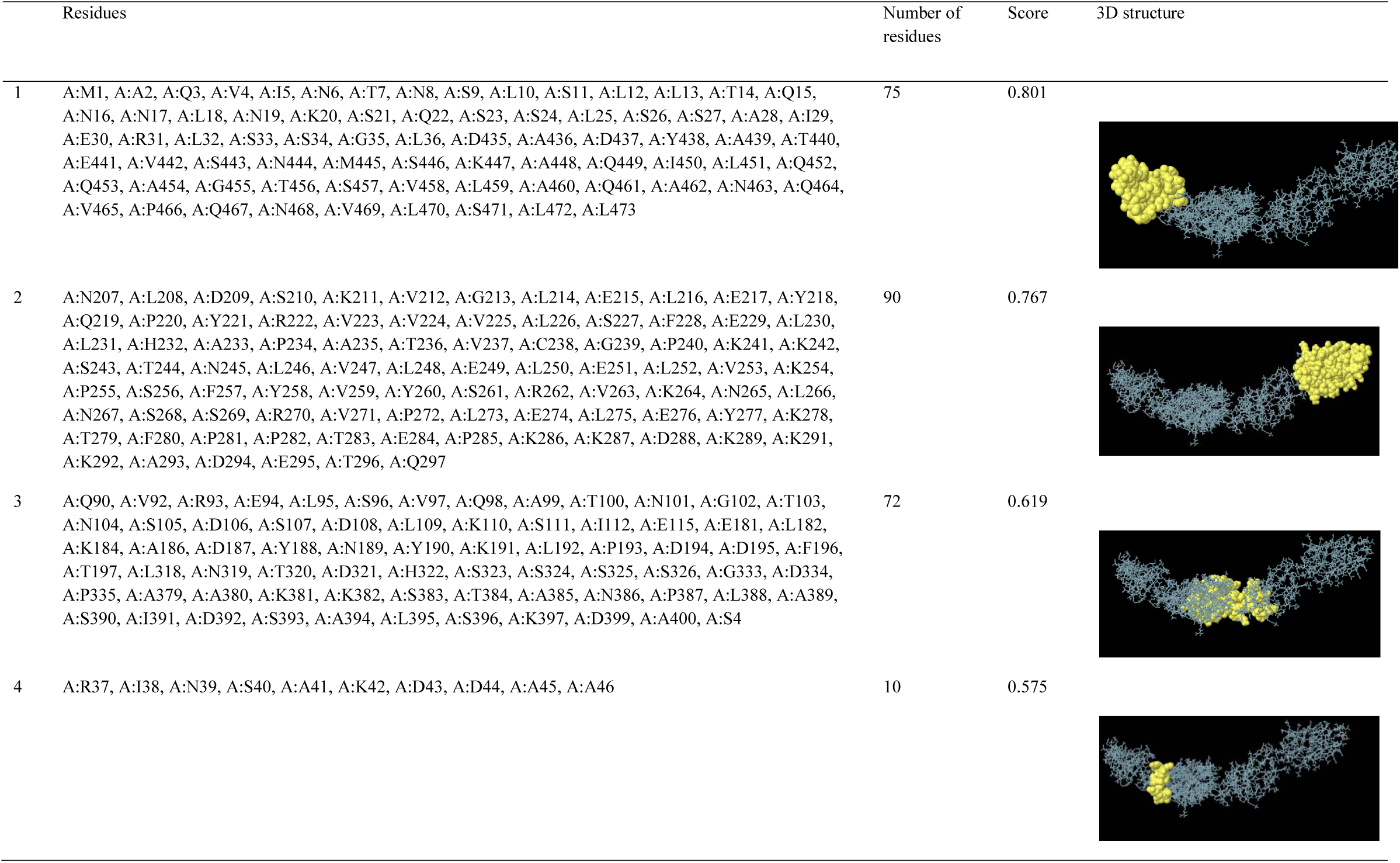
Conformational B-cell epitopes identified in the refined tertiary structure of the multi-epitope vaccine using Ellipro tool.

|  | Residues | Number of residues | Score | 3D structure |
| --- | --- | --- | --- | --- |
| 1 | A:M1, A:A2, A:Q3, A:V4, A:I5, A:N6, A:T7, A:N8, A:S9, A:L10, A:S11, A:L12, A:L13, A:T14, A:Q15, A:N16, A:N17, A:L18, A:N19, A:K20, A:S21, A:Q22, A:S23, A:S24, A:L25, A:S26, A:S27, A:A28, A:I29, A:E30, A:R31, A:L32, A:S33, A:S34, A:G35, A:L36, A:D435, A:A436, A:D437, A:Y438, A:A439, A:T440, A:E441, A:V442, A:S443, A:N444, A:M445, A:S446, A:K447, A:A448, A:Q449, A:I450, A:L451, A:Q452, A:Q453, A:A454, A:G455, A:T456, A:S457, A:V458, A:L459, A:A460, A:Q461, A:A462, A:N463, A:Q464, A:V465, A:P466, A:Q467, A:N468, A:V469, A:L470, A:S471, A:L472, A:L473 | 75 | 0.801 |  |
| 2 | A:N207, A:L208, A:D209, A:S210, A:K211, A:V212, A:G213, A:L214, A:E215, A:L216, A:E217, A:Y218, A:Q219, A:P220, A:Y221, A:R222, A:V223, A:V224, A:V225, A:L226, A:S227, A:F228, A:E229, A:L230, A:L231, A:H232, A:A233, A:P234, A:A235, A:T236, A:V237, A:C238, A:G239, A:P240, A:K241, A:K242, A:S243, A:T244, A:N245, A:L246, A:V247, A:L248, A:E249, A:L250, A:E251, A:L252, A:V253, A:K254, A:P255, A:S256, A:F257, A:Y258, A:V259, A:Y260, A:S261, A:R262, A:V263, A:K264, A:N265, A:L266, A:N267, A:S268, A:S269, A:R270, A:V271, A:P272, A:L273, A:E274, A:L275, A:E276, A:Y277, A:K278, A:T279, A:F280, A:P281, A:P282, A:T283, A:E284, A:P285, A:K286, A:K287, A:D288, A:K289, A:K291, A:K292, A:A293, A:D294, A:E295, A:T296, A:Q297 | 90 | 0.767 |  |
| 3 | A:Q90, A:V92, A:R93, A:E94, A:L95, A:S96, A:V97, A:Q98, A:A99, A:T100, A:N101, A:G102, A:T103, A:N104, A:S105, A:D106, A:S107, A:D108, A:L109, A:K110, A:S111, A:I112, A:E115, A:E181, A:L182, A:K184, A:A186, A:D187, A:Y188, A:N189, A:Y190, A:K191, A:L192, A:P193, A:D194, A:D195, A:F196, A:T197, A:L318, A:N319, A:T320, A:D321, A:H322, A:S323, A:S324, A:S325, A:S326, A:G333, A:D334, A:P335, A:A379, A:A380, A:K381, A:K382, A:S383, A:T384, A:A385, A:N386, A:P387, A:L388, A:A389, A:S390, A:I391, A:D392, A:S393, A:A394, A:L395, A:S396, A:K397, A:D399, A:A400, A:S4 | 72 | 0.619 |  |
| 4 | A:R37, A:I38, A:N39, A:S40, A:A41, A:K42, A:D43, A:D44, A:A45, A:A46 | 10 | 0.575 |  |

### 5.3 Tertiary structure modeling, refinement, and validation

The primary 3D model of the protein vaccine construct was predicted using I-TASSER. The server suggested the top five models for the protein vaccine with C scores ranging from −5 to +2. The C score of the selected model was 2.97, which was the highest score of C. Then, selected I-TASSER model was refined by GalaxyRefine software. The quality of the designed-vaccine was evaluated using the Ramachandran plot in the RAMPAGE server and characteristic atomic interaction in the ERRAT server. The RAMPAGE results of the final model showed that the majority of residues (89.6%) are located in the favored region and 6.4% are in allowed, and only 4% of residues are the outlier. The ERRAT result showed that the refined model has the ERRAT score 87.955 (Fig.2). The outputs of the Ramachandran plot and ERRAT indicated that the refined 3D structure is good and therefore, can be utilized as a reliable model for further evaluations.

**FIG.2:**
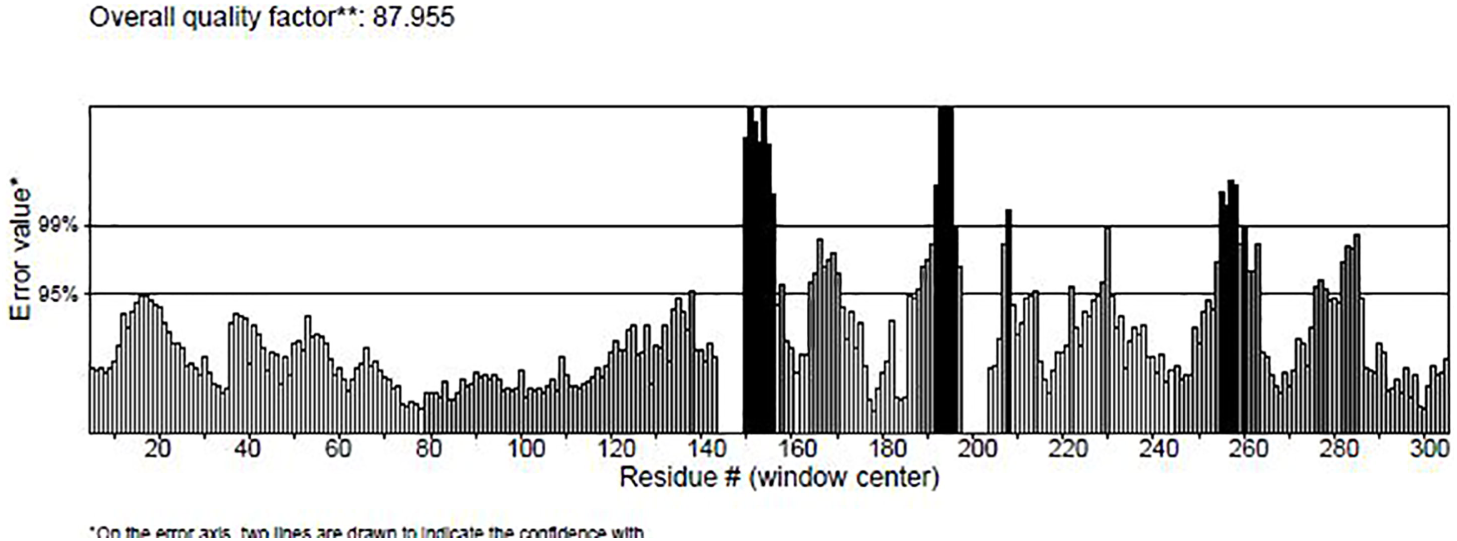
Validation results of the refined 3D structure of multi-epitope vaccine structure by ERRAT software. ERRAT plot showed the overall quality score of the refined structure as 87.955.

### 6.3 Conformational B-cell epitope Identification

Tertiary structure of the designed vaccine was used as an input for conformational (discontinuous) B cell epitope prediction via Ellipro in IEDB. From 472 amino acid residues, 247 were defined as discontinues B cell epitope.

### 7.3 Identifying binding sites and protein–protein docking

ClusPro server has been used to perform the docking of the vaccine with TLR3, 5, and 8 molecules. Cluspro generated thirty models for each of interaction. The docked model was selected based on most atoms interacted in the vaccine and each of receptors (Figures 3 to 5).

**Fig. 3.**
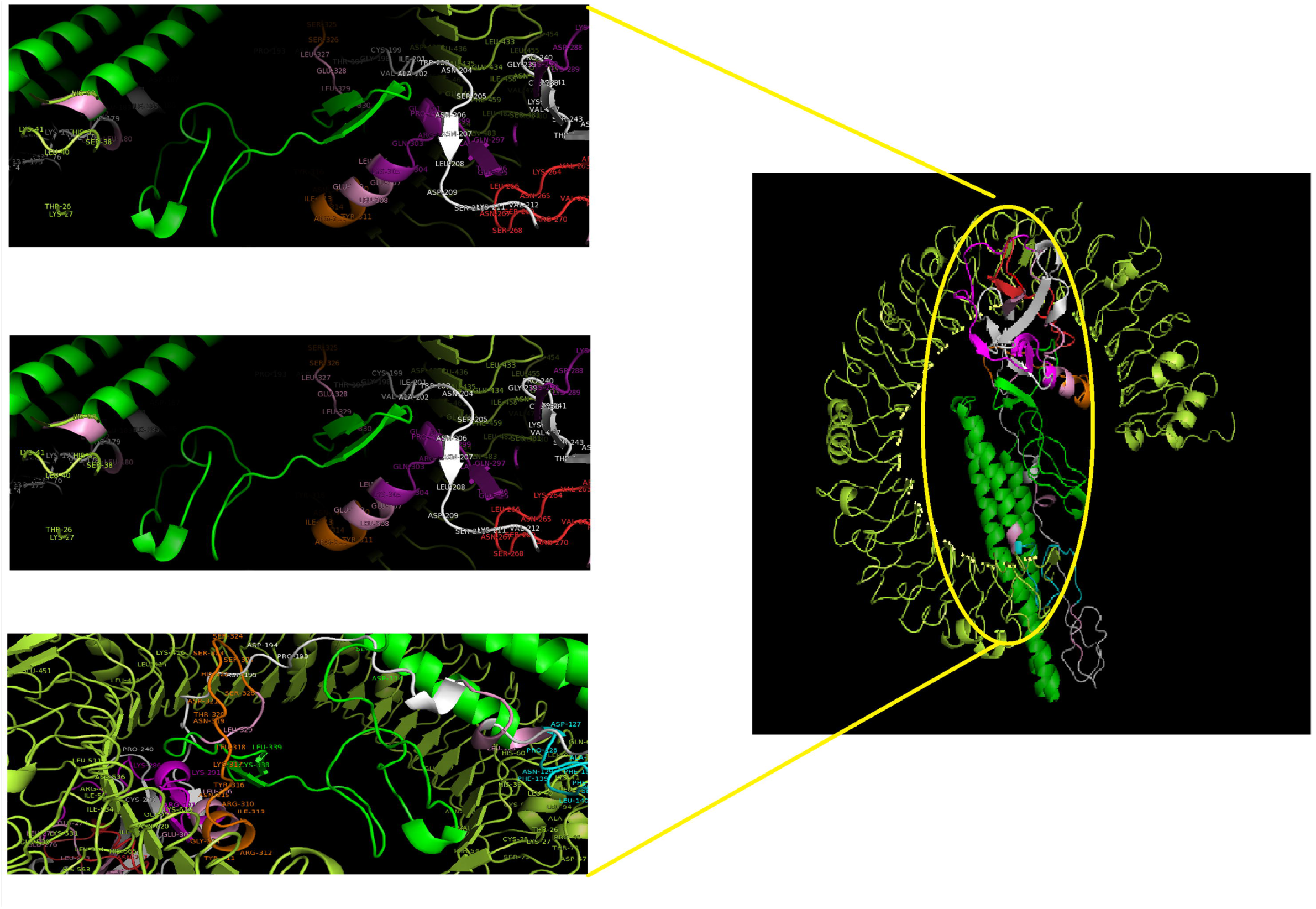
Docking model (cartoon representation) of human TLR3 in complex with the vaccine obtained using Cluspro server. TLR3 protein is shown in limon, Flagellin in green, HBD-3 in tv-green, S epitopes in white, M epitope in orange, N epitope in purple, E epitope in red, HP-91in cyan and linkers are shown in light pink. As it shows some parts of two S, M, N, and E epitopes, HBD-3, and HP-91 interacted with TLR3. To more visualized interaction points, some of the interacting residues of the vaccine and TLR3 are magnified in 20 Angstrom. Docked model was visualized via Pymol software.

**Fig. 4.**
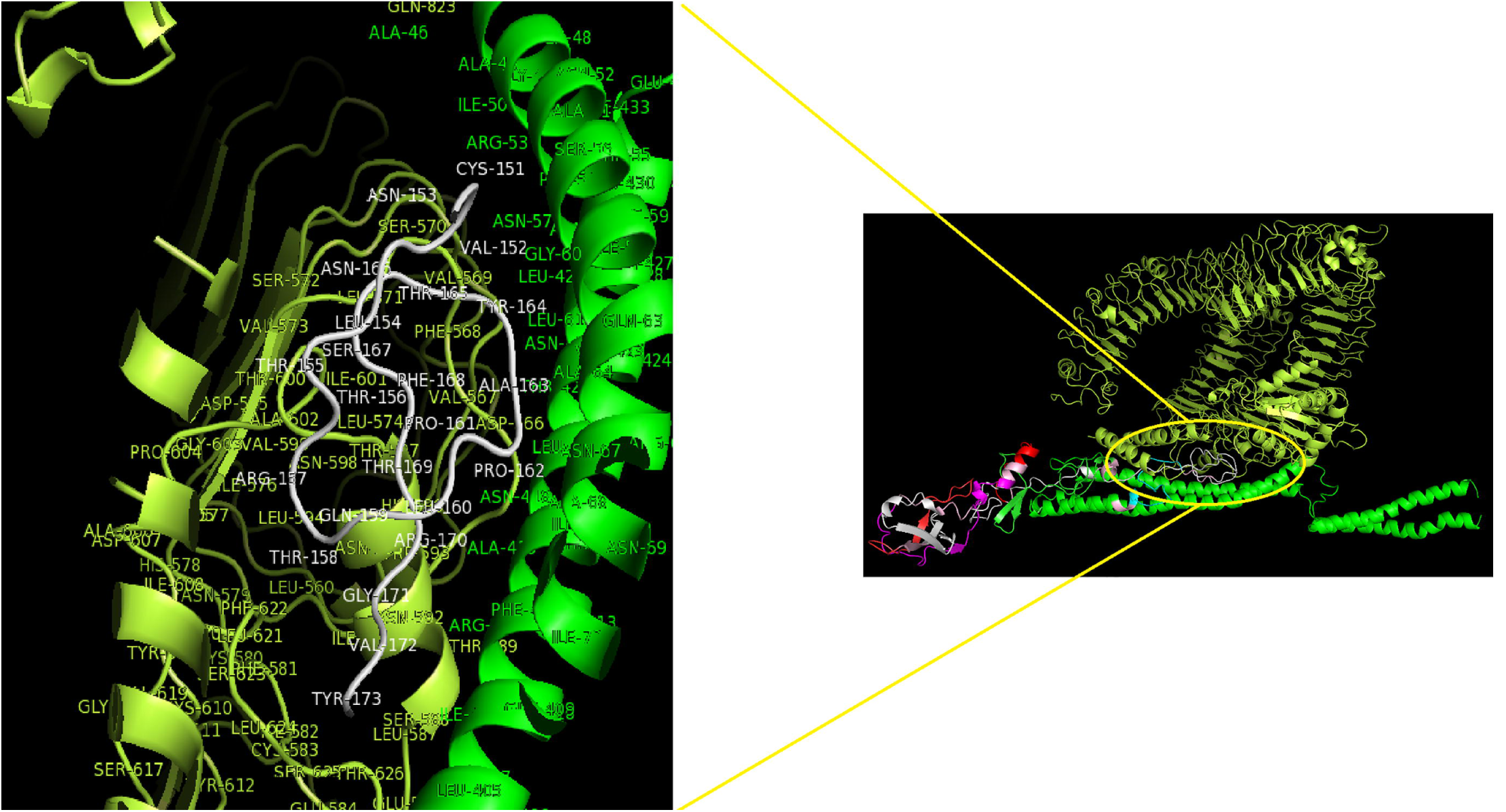
Docking model (cartoon representation) of human TLR5 in complex with the vaccine obtained using Cluspro server. TLR5 protein is shown in limon, Flagellin in green, HBD-3 in tv-green, S epitopes in white, M epitope in orange, N epitope in purple, E epitope in red, HP-91in cyan and the linkers are shown in light pink. As the figure shows some parts of Flagellin, S-epitope and HP-91 interacted with TLR5. In order to more visualized interaction points, some of the interacting residues of the vaccine and TLR5 are magnified in 20 Angstrom. Docked model was visualized by Pymol software.

**Fig. 5.**
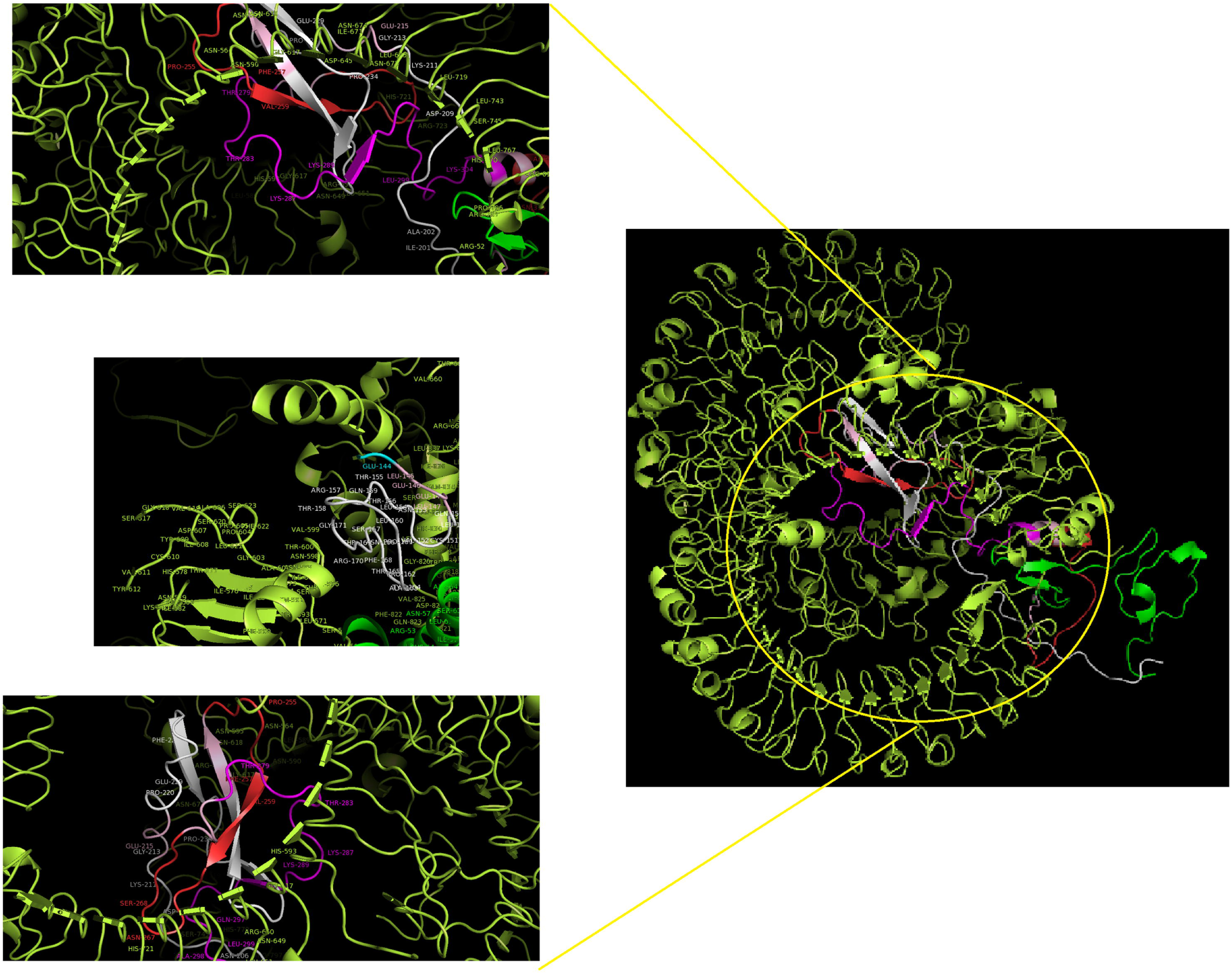
Docking model (cartoon representation) of human TLR8 in complex with the vaccine obtained using Cluspro server. TLR8 protein is shown in limon, Flagellin in green, HBD-3 in tv-green, S epitopes in white, M epitope in orange, N epitope in purple, E epitope in red, HP-91in cyan, and linkers are shown in light pink. As the figure shows some parts of S, M, N, and E epitopes, HBD-3 and HP-91 interacted with TLR8. In order to more visualized interaction points, some of the interacting residues of the vaccine and TLR8 are magnified in 20 Angstrom. Docked model was visualized using Pymol software.

### 8.3 *In silico* codon optimisation of the vaccine construct

The reverse translation of the protein vaccine into a nucleotide sequence was performed simultaneously using the Seipence Manipulation Suite server to express high-level protein in E. coli. Codon Adaptation Index (CAI), GC content and Codon Frequency and Distribution (CFD) were evaluated using GenScript online server. A CAI of the vaccine optimized nucleotide sequence was 1. The GC content of the structure was in the ideal range of 55.78% (30-70%), and CFD100 indicating the effective expression of the protein in the host. To clone the gene into *E. coli* vectors, XhoI and EcoRV restriction sites were added into the N- and C-terminals of the sequence using CLC Sequence viewer v.8. Finally, the vaccine gene was inserted into the PET-28 vector (Fig.6).

**Fig.6.**
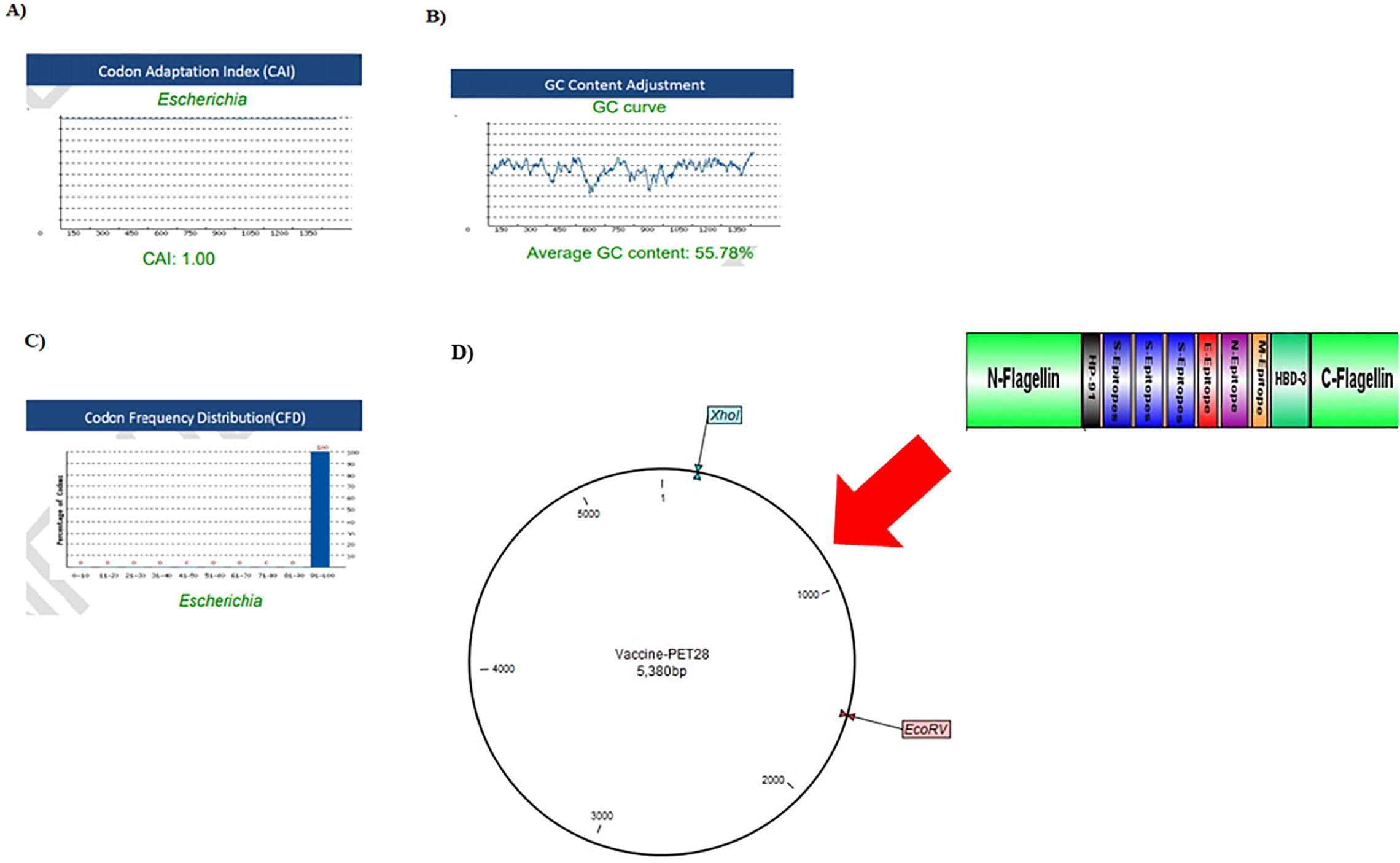
Evaluation of the three important parameters of the codon-optimized gene to express high level protein in E.coli. A) CAI of the gene sequence was 1. It is noted that a CAI of > 0.8 will be considered as good for expression in the selected host. B) Average GC content of the sequence was 55.78%. C) Codon frequency distribution (CFD) value of the gene sequence is 100. A CFD equal to100 supports maximum protein expression in the desired host. D) Insertion of the vaccine gene in the PET28 vector by restriction enzymes of XhoI and EcoRV.

## 4. Discussion

To date, there remains an utmost need to design efficient vaccines in order to halt the inexorable spread of COVID19. In light of recent advances in bioinformatics approaches, we can identify immunodominant T and B cell epitopes and design a potential vaccine to prevent the disease. We used an in silico analysis to design a potent multi-epitope peptide vaccine against SARS-CoV-2. The vaccine contains 472 amino acids which constructed from three specific epitopes from S (receptor-binding domain) and one epitope from each of the structural proteins of E, M, and N. These proteins have essential roles in infection of host cells and host immune modulation. Therefore, it is expected that the vaccine will have a high ability to induce the production of neutralizing antibodies by B lymphocytes and the production of IFN-γ by (T helper) Th and CTL cells.

Adjuvants are essential components of multi-epitope vaccines, because they increase immunologic properties of the vaccine structures. We used Flagellin, a TLR5 agonist, in two N and C terminous of the vaccine construct. Flagellin induces innate immune effectors such as cytokines and nitric oxide[70, 71], activate of TLR5-positive DCs, neutrophils and epithelial cells[72-75] and stimulate the activation of adaptive immune responses mainly Th2-type and IgA production [72, 73, 76-79]. Intranasal administration of Flagellin stimulates the signaling of TLR5 in lung epithelial cells and pneumonocytes [80]. It was used broadly as adjuvant in vaccine structure designed against viral infections such as influenza [79, 81, 82] and HPV[23, 83]. In addition, we incorporated HP-91 and HBD-3 in the final vaccine construct. The peptide of HP-91 derived from B-box domain amino acids of 89-108 from HMGB1 protein. This peptide induces high levels of IL-2 and IL-15, increases secretion of IL-12 and IFN-α in human DCs and augmented T cell activation[84]. Also, It activates (Dendritic cells) DCs independent of TLR2, 4, and 9, and MyD88 pathways[85].This is a potent immune adjuvant for inducing cellular and humoral immune responses and used in vaccines against viral infection such as HIV and HPV [84-87]. HBD-3 is the third adjuvant in our vaccine, used as adjuvant to viruses such as influenza and MERS-CoV [26, 88]. This peptide blocks viral fusion using creating a protective barricade of immobilized surface proteins[88]. It activates APCs via TLR1 and TLR2[89] and induces IL-22[90], TGF-α[91, 92] and IFN-γ[93, 94]. It has been implicated in the chemotaxis of immature DCs and T cells via its interaction with chemokine receptor 6 (CCR6) and the chemotaxis of monocytes via interaction with CCR2[95], As well as this peptide activates NK cells, promotes and activates myeloid DCs directly and dependent Natural killer (NK) activity[89, 94]. For linking the different pieces of the vaccine, a repeated of a dipeptide linker of LE was used. This linker improves the expression of vaccine protein via decreasing the pI [96, 97].

An efficient vaccine should not only have stimulating ability immune response but also possess good physicochemical properties during production, formulation, storage and consumption. According to the results of bioinformatics predictions, the designed vaccine was stable with a stability index of 35.77 and had pI of 7.45. Conformational B cell epitopes have a central role in induction humoral responses. The accessible of a significant number of B cell epitopes in the vaccine molecule indicates the high capability of this structure in the stimulation of B lymphocytes. The binding affinity of the vaccine with immune receptor (TLR3, 5 and 8) is necessary to effectively transport vaccine protein into antigen presenting cells. The results of docking analysis showed the vaccine protein properly interact with TLR3, 5, and 8.

## 5. Conclusion

In this study, using a variety of bioinformatics methods, we developed a multi-epitope subunit vaccine against the SARS-Cov-2 virus. The results of this study showed that it could be possible to predict vaccine candidates against new emerging viral diseases such as COVID-19 with the help of reverse vaccinology. The present study, with its effective vaccine design against SARS-CoV-2, showed that bioinformatics approaches could help to develop effective treatments for other emerging infectious diseases in a short time and at low cost. However, the *in silico* results of this study need to be verified using laboratory and animal models.

## 6. List of abbreviations

ACE2: angiotensin-converting enzyme 2
ANN: artificial neural network
CAI: Codon Adaptation Index
CASP10: Critical assessment of techniques for protein structure prediction
CCR: C-C Motif Chemokine Receptor
CFD: Codon Frequency and Distribution
CombLib: Combinatorial Peptide Libraries
CTL: Cytotoxic T lymphocytes
DC: Dendritic cells
E: envelope
E.coli: Escherichia coli
h: hour
HBD-3: human beta defensin 3
HMGB1: high mobility group box1
HLA: Human leukocyte antigen
KDa: Kilo Dalton
INF: Interferon
M: membrane
MHC: Major histocompatibility complex class
MW: molecular weight
N: nucleocapsid
NK: Natural killer
pI: isoelectric point
PSSMs: position specific scoring matrices
QM: Quantitative matrix
S: spike
SARS-CoV-2: severe acute respiratory syndrome coronavirus 2
SMM: stabilized matrix method
SVM: support vector machine
Th: T helper
TLR: Toll-Like Receptor
3D: Three dimensional

## 8. Declarations

### 7.2 Authors’ contributions

A.R conceived and designed this study; supervised the project and collected the data. Z.Y and M.Y contributed to performing the analysis and writing the draft. A.R and R.V edited the manuscript.

### 7.3 Ethics approval and consent to participate

The protocol of this study approved by Research Ethics committee of Mazandaran University of Medical Sciences IR.MAZUMS.REC.1398.1430.

### 7.4 Consent for publication

Not applicable.

---

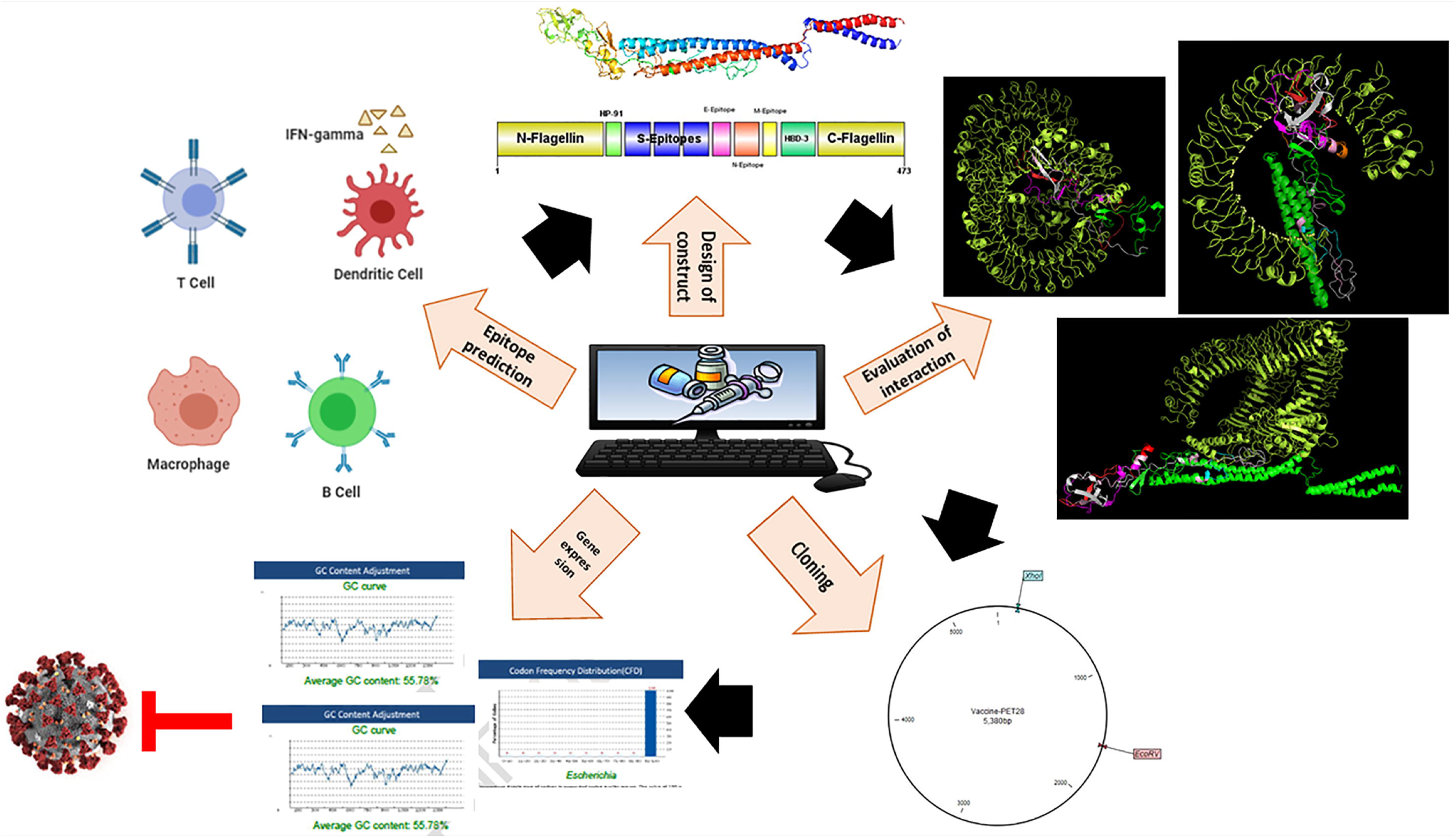

